# A modular architecture for trial-by-trial learning of redundant muscle activity patterns in novel sensorimotor tasks

**DOI:** 10.1101/2025.01.29.635432

**Authors:** Lucas Rebelo Dal’Bello, Denise Jennifer Berger, Daniele Borzelli, Etienne Burdet, Andrea d’Avella

**Affiliations:** Laboratory of Neuromotor Physiology, Fondazione Santa Lucia IRCCS, Rome, Italy; Department of Systems Medicine and Centre of Space Bio-medicine, University of Rome Tor Vergata, Rome, Italy; Department of Biomedical and Dental Sciences and Morphofunctional Imaging, University of Messina, Messina, Italy; Department of Bioengineering, Imperial College of Science, Technology and Medicine, London, UK; Department of Biology, University of Rome Tor Vergata, Rome, Italy

**Author notes:** Corresponding author: (LD).

## Abstract

The coordination of the multiple degrees-of-freedom of the human body may be simplified by muscle synergies, motor modules which can be flexibly combined to achieve various goals. Studies investigating adaptation to novel relationships between muscle activity and task outcomes found that altering the recruitment of such modules is faster than the learning of their structures *de novo*. However, how learning new synergy recruitments or new synergy structures may occur remains unclear. While trial-by-trial learning of novel sensorimotor tasks has been successfully modeled at the level of task variables, few models accounted for the redundancy of the motor system, particularly at the muscular level. However, these models either did not consider a modular architecture of the motor system, or assumed *a priori* knowledge of the sensorimotor task. Here, we present a computational model for the generation of redundant muscle activity where explicitly defined modules, implemented as spatial muscle synergies, can be updated together with their recruitment coefficients through an error-based learning process dependent on a forward model of the sensorimotor task, which is not assumed to be known a priori. Our model can qualitatively reproduce the experimental observations of slower learning and larger changes in the structure of the muscle activity under sensorimotor tasks that require the learning of novel patterns of muscle activity, providing further insights into the modular organization of the human motor system.

**Author summary:** It has been proposed that muscles are recruited in modules, called muscle synergies, rather than on a muscle-by-muscle basis. This modular organization was shown to affect the learning of novel tasks, where simulated remapping of the forces generated by the muscles (“virtual surgeries”) that make the synergies ineffective are more difficult to learn. Previous models of trial-by-trial learning in a modular architecture have assumed prior knowledge of the motor task (in particular, how to modify the muscles / synergies recruitment given an error), which might not be the case for tasks in which novel patterns of muscle activity are required. In contrast, models which assumed no prior knowledge of the task have not investigated the role of modularity. Here, we propose a computational model of the trial-by-trial adaptation of muscle synergies, their recruitment, and the concurrent learning of the internal model of the musculoskeletal system responsible for error correction. Our results show that this model replicates experimental observations of slower learning and larger changes in the structure of the muscle activity in sensorimotor tasks that require the learning of novel patterns of muscle activity, providing insight into learning-related changes of muscle activity in novel sensorimotor tasks.

## 1 Introduction

During movement planning and execution, the central nervous system needs to coordinate the activity of a large and redundant set of muscles acting on multiple joints [1]. It has been suggested that the problem of coordinating the multiple degrees-of-freedom of the human body is simplified by grouping multiple muscles into a reduced number of motor modules, often called muscle synergies, which can be flexibly combined to generate a large repertoire of movements [2]. The modularity of the motor system has been investigated through the decomposition of the muscle activity patterns measured either during voluntary movements or in response to cutaneous or spinal cord electrical stimulation. The key observation is that the electromyography (EMG) signals recorded from many muscles can be accurately reconstructed by the combination of a small number of modules which generalize across multiple contexts and movements [3–6]. However, this low-dimensionality observed in the muscle activity patterns may be due to task and biomechanical constraints rather than to a modular organization of the motor system [7].

Further support for the modular organization of the motor system has come from testing specific predictions of how modularity would affect a novel motor learning task [8]. In this task, participants generated a virtual force to control a cursor using their isometric muscle activity measured with EMG and adapted to two different perturbations of the mapping of muscle activity to virtual force, or “virtual surgeries”. These perturbations consisted of changes to the directions of the virtual forces generated by the contraction of each muscle, akin to the effect of a complex tendon-transfer surgery involving multiple muscles. The two types of surgeries differed in how effective the muscle synergies identified before the surgery were in generating a virtual force. After compatible surgeries the muscle synergies still spanned the entire force space, while incompatible surgeries reduced the space of forces that can be spanned by the muscle synergies. However, the force space was still spanned by the individual muscles under both types of surgeries, so that a modular organization of the motor system predicted a difference in the adaptation rate to the two types of surgeries, while a non-modular organization predicted no difference. The observation of a significantly slower learning after incompatible virtual surgeries provided stronger evidence to the modular organization of the motor system.

In addition to their role on movement generation, muscle synergies also play a role in motor adaptation and motor learning. During visuomotor rotations, it has been shown that the directional tuning of the synergy recruitment is a possible adaptation strategy employed by the central nervous system [9]. As previously mentioned, in studies that investigated the adaptation to novel relationships between muscle activity and the resulting end-effector force using virtual surgeries, it was shown that surgeries which were compatible with the original synergies (requiring only a change in the synergy directional tuning) were learned faster than surgeries which were incompatible (requiring the usage of novel muscle activity patterns), suggesting that the altering of the synergy recruitment coefficients is faster than the learning *de novo* of new synergy structures [8,10]. These results suggest that the modularity of the motor system influences its adaptation to perturbations and its learning of novel patterns of activity, which may be relevant to neurorehabilitation [11].

While the adaptation of hand reaching movements under externally-induced visuomotor errors has been extensively studied and modeled [12], relatively few studies have focused on changes at the highly-redundant muscular level. The adaptation of feedforward muscle activity in response to feedback error under force-field perturbations has been modeled by a “V-shaped” learning function, explaining the regulation of reciprocal- and co-activation in stable and unstable conditions [13]. Similar to changes to the directional tuning of muscle synergies under visuomotor rotations [9], changes in muscles’ tuning curves under force-field perturbations have also been reported [14]. When accounting for the modularity of the motor system, the adaptation to virtual surgeries incompatible with the original synergies produced muscle activity patterns poorly explained by the original synergies [8]. Such change in the muscle activity patterns was shown not to interfere with performance in the baseline task, and was persistent even during a subsequent exposure to a compatible surgery, suggesting an increase in the exploration of muscular null-space activity patterns [10]. Overall, however, the error-based learning mechanisms governing changes at the muscular level, including learning of novel muscle activity patterns, are still unclear.

Learning of novel patterns of muscle activity, and not simply reusing already known patterns, is linked to the acquisition of new motor skills, a process also known as *de novo* learning [15]. While the exact mechanisms of *de novo* learning are still unclear, it has been argued that it requires the learning of an internal model responsible for error corrections, which can then be used to guide the update of the controller for the motor task [16–18]. A few studies have investigated the learning of such error-correcting internal models simultaneously through a trial-by-trial update of a feedforward controller [19], with one study focusing on the role of motor exploration on the learning process [20]. However, these studies did not account for the possible modularity of the motor system. In contrary, studies that include modularity assumed that the error-correcting internal model was known a *priori*, which might not be the case in *de novo* learning [21,22]. Also, one of these studies [22] did not investigate a possible update of the synergies’ structure, which has been observed during training under incompatible virtual surgeries [8,10].

We introduce here a model of trial-by-trial, error-based learning of redundant motor commands in a modular control architecture to address these open issues of computational models of learning in the redundant motor system. We model the feedforward generation of redundant muscle activity patterns during isometric force-reaching tasks through explicitly defined muscle synergies, and the learning of novel motor commands as the trial-by-trial update of both muscle synergies and their recruitment coefficients at different learning rates. These updates are performed by the backpropagation of the difference between the intended and executed force through a forward model of the isometric task. Importantly, this forward model is also updated over time, reflecting learned changes in the sensorimotor system or in the task environment. We show how our model can qualitatively reproduce the results of tasks with multidirectional isometric force generation under different perturbations, such as visuomotor rotations and virtual surgeries that rearrange the directions of the forces produced by the muscles. We analyze the predictions of our model in terms of changes in the force error and in the structure of the muscle activity across the different perturbations and the different combinations of model parameters that were simulated.

## 2 Methods

### 2.1 Computational model of trial-by-trial adaptation

We model the trial-by-trial adaptation of the redundant muscle activity patterns during the generation of isometric force at the hand (Fig 1) in a musculoskeletal system that can be exposed to perturbations such as visuomotor rotations and ‘virtual surgeries’ [8]. In our model, we assume that the generation of the redundant muscle activity occurs through the recruitment of explicitly defined modules representing spatial muscle synergies. Then, a muscle pattern ***m***, i.e., a non-negative *M*-dimensional vector of activation of a set of *M* muscles, is generated by the linear combination of *N* muscle synergies, each a non-negative *M*-dimensional vector:

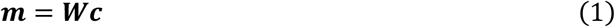

**Fig 1:**
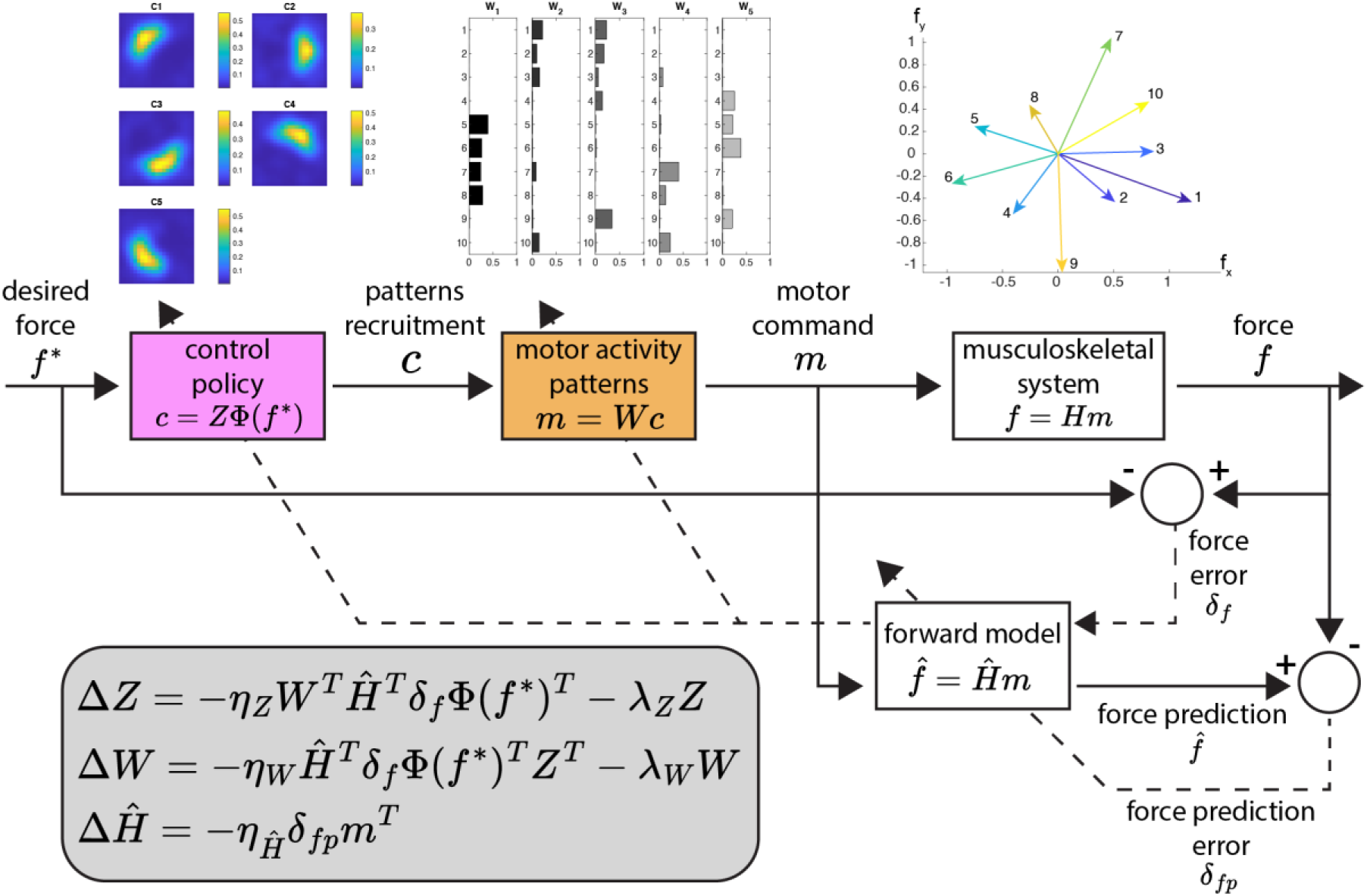
Computational model of trial-by-trial generation and update of redundant muscle activity for isometric force control. The model maps a desired force into a recruitment of muscle synergies using a radial basis function-based control policy. The combination of the muscle synergies results in the muscle activity executed through the musculoskeletal system. Both the control policy and the synergy structure can be updated by backpropagating the force error through a forward model of the musculoskeletal system. This forward model can also be updated using force prediction error.

where ***W*** is an *M* × *N* matrix with the synergy vectors as columns, and ***c*** is a *N*-dimensional synergy combination vector. The activation of each muscle generates a specific isometric force at the virtual hand which we approximate as a linear function of activation, and the activation of all muscles when added result in an end-effector force ***f*** (*D*-dimensional vector):

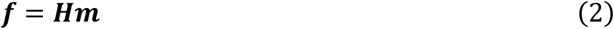

where ***H*** is the *D* × *M* matrix that linearly maps the muscle activation pattern into a *D*-dimensional force.

The synergies are recruited according to a control policy that maps a desired force ***f***^∗^ into synergy recruitment coefficients. We implement this control policy using radial basis functions, which have been shown to reproduce well the generalization patterns observed in motor adaptation experiments and have been extensively used in fitting primitives of internal models for movement planning [23–25]. We implement this radial basis function-based control policy with:

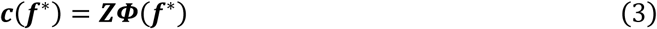

where ***Z*** is a *N* × *N_Φ_* matrix of combination coefficients and ***Φ***(***f***^∗^) is a *N_Φ_*-dimensional vector of Gaussian basis functions activations, with centers spread in a region of the *D*-dimensional force space. The generation of the muscle activity can then be written as:

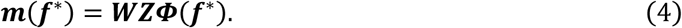

In our simulations, we also add signal-dependent, random motor noise to the muscle activity obtained from the model, sampled from a multivariate Gaussian distribution with the variance of each muscle scaled quadratically with that muscle’s activation [26] and with zero covariance between different muscles.

In contrast to other models of the feedforward generation of redundant muscle activity [21,22], which assumed that the knowledge of the environment (that is, the relationship between muscle activation and the resulting force visualized at the end-effector, which includes the musculoskeletal system and possible perturbations of the resulting force) is known *a priori* (or that it is learned instantly when the environment is perturbed), in our model the environment must be learned by practice, and is represented by an internal estimation ***Ĥ*** of the muscle activity-to-force matrix ***H***. This estimation ***Ĥ*** can be used as an internal forward model to predict forces ***f̂*** that are generated given a muscle activity ***m***:

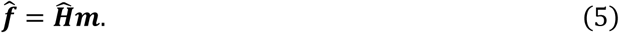

In our model, a control policy is learned by updating the synergy structure matrix ***W*** and the combination coefficients matrix ***Z***. The goal of the learning process is to minimize the squared norm of the force error ***δ***_*f*_ = ***f*** − ***f***^∗^ defined as the difference between the executed force ***f*** and the desired force ***f***^∗^. The matrices ***W*** and ***Z*** are then updated based on the gradient of the squared norm of the error, 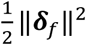, with respect to each matrix (derivations of the update equations are shown in S1 Text). We also add a regularization term to each matrix update to reduce co-contraction of muscles and co-recruitment of synergies, i.e. the effort.

For the synergy structure matrix ***W***, the update *Δ****W*** after a trial with a desired force ***f***^∗^ where an error ***δ***_*f*_ is observed becomes:

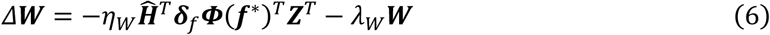

where *η*_*W*_ is a scalar learning rate and *λ*_*W*_ is a scalar regularization weight. This is in contrast with previous models of the generation of redundant motor activity where the synergies were considered fixed [22], or where a modular controller was not considered altogether [19,20].

The update *Δ****Z*** of the combination coefficients matrix ***Z*** becomes:

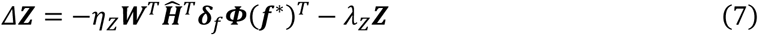

where *η*_*Z*_ is a scalar learning rate and *λ*_*Z*_ is a scalar regularization weight. It is important to note that both update equations contain the matrix ***Ĥ***^*T*^ of the internal forward model of the environment, which here acts as an estimation of the “sensitivity derivative” 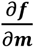 of the environment [27]. When calculating the gradient of the squared norm of the error 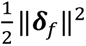, both update equations would contain the matrix *H*^*T*^ of the real environment which is not *a-priori* known. Since we assume that the learner uses an internal estimation of the environment, the matrix ***Ĥ***^*T*^ is used instead in the update equations [16].

Equations 6 and 7 suggest that if there is a large discrepancy between the environment ***H*** and the internal model of the environment ***Ĥ***, the update of matrices ***W*** and ***Z*** of the controller will not necessarily minimize the norm of the error ***δ***_***f***_. Indeed, it has been shown that a sufficient condition for the internal model of the environment ***Ĥ*** to decrease the squared norm of the error during the update of the model is that 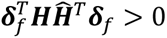, that is, the vector ***Ĥ***^*T*^***δ***_*f*_ (present in both update equations) has an angle of less than 90° with the vector ***H***^*T*^***δ***_*f*_, which uses the original environment matrix ***H*** [28].

We update the internal forward model ***Ĥ*** by using the gradient of the norm of the prediction error ***δ***_*fp*_ = ***f̂*** − ***f*** between the force predicted by the forward model given a muscle activity ***m*** and the actual generated force for the same muscle activity. The update *Δ****Ĥ*** can be written as:

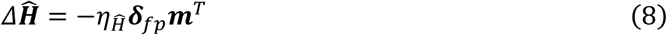

where *η*_*Ĥ*_ is a scalar learning rate.

### 2.2 Model simulations

We performed simulations to validate the model by assessing whether it can adapt to perturbations such as visuomotor rotations and ‘virtual surgeries’ and predict key features of motor adaptation experiments with human participants. All our simulations and analyses were conducted in MATLAB^Ⓡ^ (MathWorks Inc., Natick, MA). In our simulations, we selected the arbitrary numbers of *D* = 2 dimensions of force, *M* = 10 muscles, and *N* = 5 muscle synergies, values similar in magnitude to the dimension of the muscle activity and number of synergies acquired during experiments with humans [8].. The muscle activation-to-force matrix ***H*** was initialized by assigning random *D*-dimensional forces to each of the *M* muscles, ensuring that the forces positively spanned the entire force space [29], and randomly assigning the norm of each vector to be in the range between 0.5 and 1.5 (arbitrary units). We defined 8 target forces ***f***^∗^, uniformly spread in a circle distant 0.5 from the center of the force space (at rest), and we initialized the *N*_***Φ***_ = 121 Gaussian basis functions ***Φ*** with centers uniformly distributed between the ranges of −1 and +1 across the two dimensions of the force space, in a 11-by-11 grid, enough for the basis functions to adequately represent all possible force targets. Each element of the matrix ***Z*** of the control policy was initialized randomly with a uniform distribution on the interval [0,0.05]. For the initialization of the matrix ***W*** of the muscle synergies, we first calculated *N* random force vectors positively spanning the entire force space [29], and then quadratic programming (MATLAB^Ⓡ^ function *quadprog*) was used to find the minimum norm, non-negative muscle activity that, in the given environment matrix ***H***, generates each of the *N* force vectors. These muscle activity vectors (each representing a muscle synergy) were concatenated, resulting in the muscle synergies matrix ***W***.

Our model was trained under three different types of perturbation: visuomotor rotations, compatible virtual surgeries, and incompatible virtual surgeries (Fig 2A). Visuomotor rotations applied a 45° counter-clockwise rotation to the force ***f*** executed by the model. Compatible and incompatible virtual surgeries correspond to changes in the pulling directions of the muscles in the musculoskeletal system, defined in terms of whether the generation of force is compatible or not with the model’s muscle synergies. They were initialized using the procedure described in [8], utilizing the environment matrix ***H*** and the muscle synergies matrix ***W*** of the model at the start of the simulation to calculate the bases of the subspaces necessary for the rotations in the muscle space (namely the basis ***N***__*c*__ of the common subspace between synergies and the null space of ***H***, the basis ***W***_**n**_*c*__ of the synergy vectors not in the null space, and the basis ***N***_**n**_*c*__ of the null space vectors not generated by synergy combinations) and calculating the compatible rotation matrix ***T***__*c*__, that rotates a vector ***w*** in the span of ***W***_**n**_*c*__ onto a second vector ***w***′ in the same subspace, and the incompatible rotation matrix ***T***_**i**_, that rotates a vector ***w*** in the span of ***W***_**n**_*c*__ onto a vector ***n*** in the span of ***N***_**n**_*c*__. The angle of the compatible rotation was adjusted so that both compatible and incompatible virtual surgeries would have a similar “index of difficulty”, defined as the average change in the muscle activity across muscles and force targets required to perform the task after the surgeries [8]. The effects of the virtual surgeries on baseline muscle synergies during representative simulations can be seen in Fig 2A, where it can be seen that compatible surgeries do not require changes to the muscle synergies in the model, but only changes to their relative recruitment along each force direction, as the forces generated by the synergies (*black arrows*) after the perturbation are altered but still span the force space. In contrast, incompatible surgeries require learning of new patterns of muscle activity, as the perturbed synergy forces are aligned in a single direction and do not span the force space.

**Fig 2:**
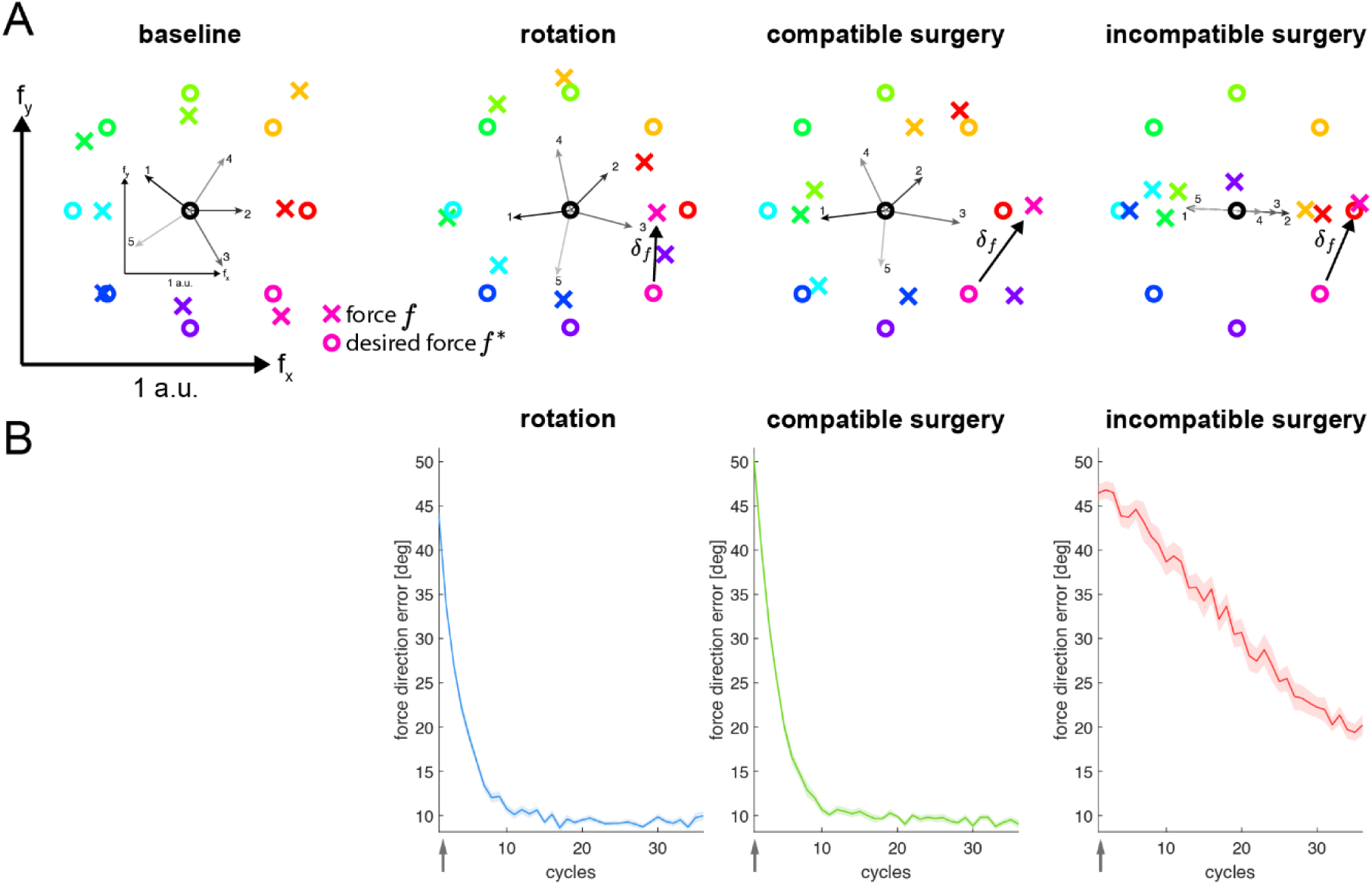
Simulated perturbations and qualitative reproduction of experimental results. (A) Diagrams show the force generated by the computational model (*colored crosses*) to match each of 8 targets (*colored circles*) at baseline and at the onset of each of three types of perturbation (rotation, compatible surgery, incompatible surgery), as well as the effect of the perturbations on the forces generated by the five muscle synergies (*black arrows*, numbered and shown in a smaller scale). During baseline, all force targets can be reached adequately (small errors are mostly due to motor noise). During a visuomotor rotation, the generated forces are rotated around the center of the task space, resulting in force errors. During a compatible surgery, the synergy forces are altered but still span the entire force space. In contrast, during an incompatible surgery, the baseline muscle synergies do not span the entire force space and the generated forces are initially all aligned along a single axis. (B) Force direction error of the simulated model during adaptation to the three perturbations (average across sixteen initializations of the model).

Each simulated experiment started with the initialization of the model and with a baseline training without any perturbation for a total of 324 cycles, with a cycle consisting of eight trials of reaching towards each of the eight defined targets in random order. During each trial, the learning rules described above were applied to update the matrices ***Z***, ***W***, and ***Ĥ*** of the model. After the initialization, each simulated experiment was trained in the baseline condition for 36 cycles, followed by 36 cycles of perturbation (the same number of trials as in [8]), and finally by 36 cycles of washout (again in the baseline condition). Within one model initialization, the data at each trial was averaged across four repetitions of the simulation, in order to attenuate the effect of the motor noise added to the motor commands at every trial. All the reported results (except individual reaching trials) were obtained by averaging at every cycle the data across 16 random initializations of the model, each with their own environment matrix ***H*** and muscle synergies ***W*** used for force generation (because of this, each initialization also had different pairs of compatible and incompatible virtual surgeries), in order to mimic an experiment with multiple participants (for comparison, the virtual surgeries results reported in [8] were obtained from 8 human participants).

We conducted three simulations under different combinations of parameters of the model, namely the learning rates of the control policy *η*_*Z*_, of the muscle synergies *η*_*W*_, of the forward model *η*_*Ĥ*_, and the regularization weights of the control policy *λ*_*Z*_ and of the muscle synergies *λ*_*W*_.

Simulation 1 tested the overall model and in particular the relative adaptation of and with synergies. To this purpose we set two possible values for the learning rate of the control policy *η*_*Z*_ and of the muscle synergies *η*_*W*_, resulting in three combinations of parameters: {(0.05,0), (0.05,0.05), (0,0.05)}. The regularization weights of the control policy, *λ*_*Z*_, and of the muscle synergies, *λ*_*W*_, were set to 1% of the learning rate of each respective component, which with a learning rate of 0.05 corresponds to a value of 5 ⋅ 10^−4^, with the same order of magnitude as used in other simulation studies [21]. The learning rate of the forward model *η*_*Ĥ*_ was fixed at 0.25. This value was chosen, along with the combination (0.05,0.05) for the learning rates of the control policy and of the muscle synergies, respectively, based on an initial sensitivity analysis where we investigated 5 different values for each parameter across 8 different model initializations, and this combination was chosen based on how well the results of the simulations qualitatively reproduced the experimental results of [8] in terms of differences, between compatible and incompatible virtual surgeries, in force direction error and in reconstruction quality of the muscle activity using the original muscle synergies. With this simulation, we could examine the relative contribution of the update of each of the components of the model in the overall learning process. All combinations of parameters were repeated across the 16 different initializations of the model.

In simulation 2, we investigated the effect of the learning of the forward model ***Ĥ*** in the overall learning process. We conducted simulations with two different learning rates *η*_*Ĥ*_, in {0,0.05}, and we also investigated a third possibility, which is that the forward model during the perturbation is an “ideal” forward model of the environment, that is, it is equal to the perturbed environment matrix ***H*** (and likewise equal to the unperturbed environment matrix during the baseline and washout periods), which corresponds to an instantaneous learning of the forward model or *a priori* knowledge of the environment as was done in previous studies [21,22]. In this simulation, the learning rates of the control policy *η*_*Z*_ and of the muscle synergies *η*_*W*_ were both fixed at 0.05, and like before the regularization weights of each component were set to 1% of their respective learning rates.

In simulation 3, we investigated the effect of the regularization in the learning process. The learning rates of the control policy *η*_*Z*_ and of the muscle synergies *η*_*W*_ were both fixed at 0.05 and the learning rate of the forward model *η*_*Ĥ*_ was fixed at 0.25. Both regularization weights *λ*_*Z*_ and *λ*_*W*_ were set at either 1% of the respective learning rate or 0.

### 2.3 Performance metrics

Across all simulations, we selected several metrics to quantify the performance of the model, calculated per each cycle of 8 trials. During each trial, the model generated force toward one of eight different force targets, with all eight targets included in a cycle presented in random order. The first two metrics that we considered, *force direction error* and the *force magnitude error*, were closely related to the performance of the reaching movement. The *force direction error* was defined as the unsigned angle (in degrees) between the line connecting the center of the force space and the force target, and the line connecting the center of the force space and the force executed by the model. The *force magnitude error* was defined as the norm of the difference between the force target and the executed force (in arbitrary units). Two additional metrics related to the muscle activity generated by the model were also calculated. The *reconstruction quality of the muscle activity using the original muscle synergies* was calculated using nonnegative linear least-squares optimization (MATLAB^Ⓡ^ function *lsqnonneg*) to find the nonnegative synergy recruitment coefficients that, when multiplied by the original muscle synergies of the simulation, minimized the squared norm of the difference with the muscle activity measured across the cycles of the simulation. We then calculated the fraction of total variation of the reconstructed muscle activity explained by the synergies (*R*^2^) across the trials of each cycle. The second metric related to the muscle activity was the *norm of the muscle activity*, averaged across the eight vectors of muscle activity generated in a cycle. In simulation 1, under the parameter combination of a non-zero learning rate for both control policy and synergies structure, the norm of the muscle activity was also evaluated in different subspaces of the muscle activity space, namely the *baseline task space* (the subspace spanned by the transpose of the baseline environment matrix ***H***), *the null space* (the subspace which results in a zero change in the force executed in the environment ***H***), the *N*_*c*_ *space* (the common subspace between the null space and the baseline muscle synergies), and the *N*_*nc*_ *space* (the subspace of the null space not intersecting the baseline muscle synergies). The norm of the muscle activity in these subspaces was calculated by projecting the muscle activity vectors in each subspace and calculating their norms.

### 2.4 Statistical analysis

We analyzed our simulated data by fitting generalized linear mixed-effects models on the data at the last cycle of the perturbation phase, using as dependent variable either force direction error, force error, fraction of total variation of the reconstructed muscle activity explained by the original synergies (*R*^2^), or norm of muscle activity (in the entire muscle activity space and in each subspace), and using as random effects the ID of each initialization of the model. In simulations 1 and 2, we used the perturbation type (rotation, compatible surgery, or incompatible surgery) the model parameter combination, and their interaction as independent variables. For simulation 1, we also fit separate models using as dependent variables the norm of the muscle activity in different subspaces, only for the model parameter combination when both control policy and muscle synergies are updated. In simulation 3, we verified that only the model for the norm of the muscle activity had a significant effect of the interaction term, so for all the other metrics we fitted models without the interaction term. Post-hoc analyses were conducted by applying F-tests to the fitted models using different contrasts, from which Bonferroni-corrected *p*-values were computed. In simulation 1, we defined the contrasts across all different perturbations and under each model parameter combination simulated (for the models with the dependent variables the norm of the muscle activity in different subspaces, contrasts were set across all different perturbations). In simulation 2 (and in the model fitted to the motor command norm in simulation 3, which included an interaction between perturbation type and model parameter combination), the contrasts were set across the different parameter combinations under each of the three perturbations.

## 3 Results

### 3.1 Qualitative reproduction of experimental results

In Fig 2B, we show the force direction error over training during simulations under different perturbations, namely a visuomotor rotation, a compatible virtual surgery, and an incompatible virtual surgery, updating all elements in the model. The force direction error during the visuomotor rotation decreases substantially in a short number of trials (each cycle corresponds to eight trials of reaching towards the eight different targets), comparable with experiments which investigated visuomotor rotations [30,31]. Importantly, the decrease in the force direction error during the incompatible surgery is much slower compared to during the compatible surgery, which is in line with experimental findings using such perturbations [8].

### 3.2 Effect of different combinations of the learning rate of the control policy and of the learning rate of the muscle synergies

For a more detailed examination of the model, we simulated the three perturbations under different combinations of the learning rates of the adaptive elements included in the model. Specifically, we investigated combinations where the control policy matrix ***Z*** and the muscle synergies matrix ***W*** have a learning rate of either zero or nonzero (the combination where both learning rates are zero was not considered). We show the results of these simulations in Fig 3. In terms of force direction error (*first row*) and force magnitude error (*second row*) during the rotation and compatible surgery perturbations, the decrease in the error appears faster when both the control policy ***Z*** and the muscle synergies ***W*** are updated (*third column*), in comparison with the simulations where only either the control policy (*first column*) or the synergies (*second column*) is updated. During the incompatible surgery the error does not decrease at all when only the control policy is updated. The error does decrease when the muscle synergies are updated, albeit slowly. Interestingly, the error decreases even faster when both control policy and synergies are updated. This is because the incompatible virtual surgeries perturb the forces generated by the muscles such that the baseline muscle synergies do not span the entire force space, therefore a change in the synergies structure is necessary to reach all directions of the force space and thus to reduce the force error. Statistical analyses on the force direction error (S1 Table) and the force magnitude error (S2 Table) indicate no significant differences between the errors (at the last cycle of the perturbation) between the rotation and the compatible surgery perturbations, but significantly larger errors for the incompatible surgery compared to the other two perturbations, under all three combinations of model parameters/learning rates.

**Fig 3:**
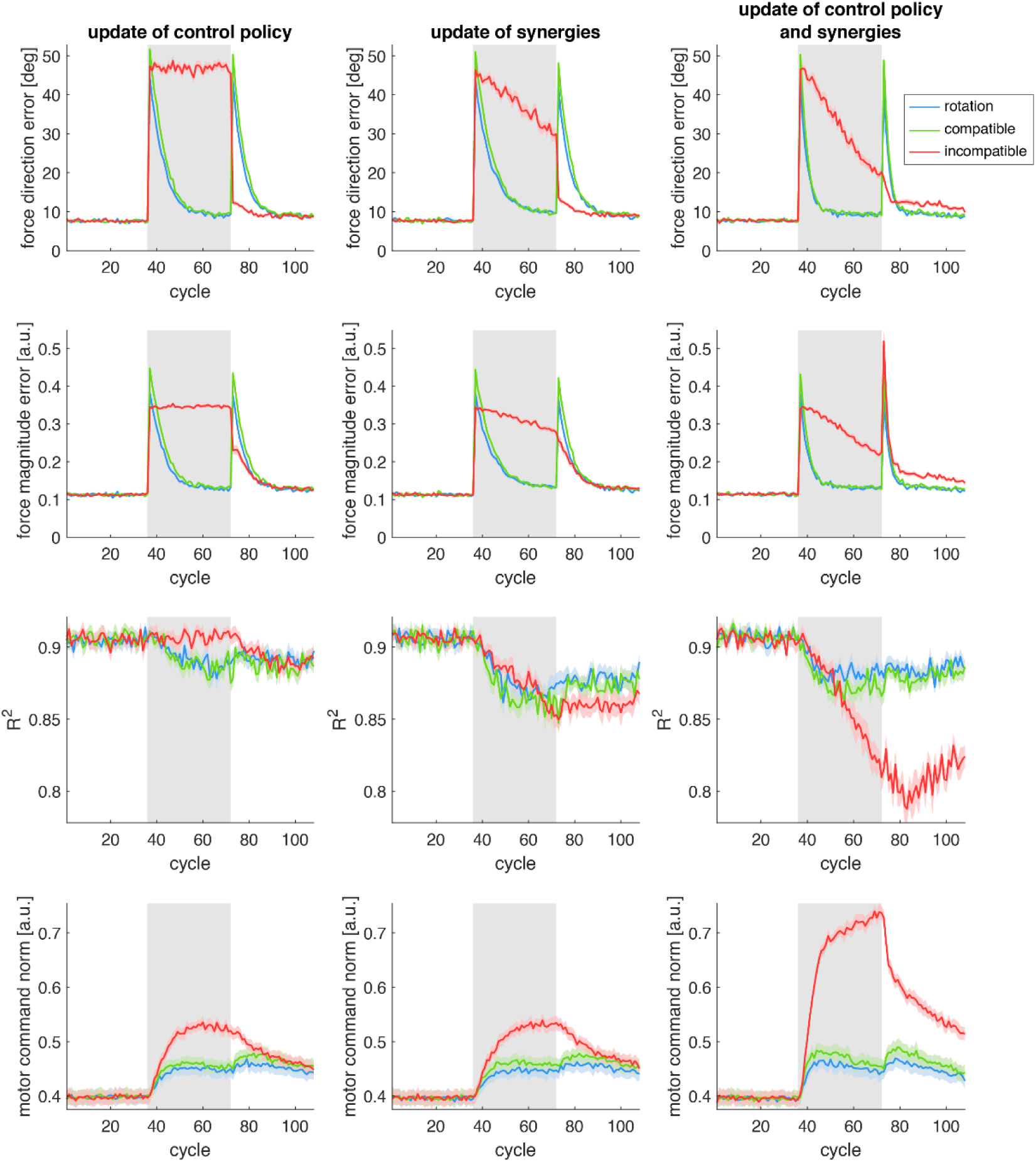
Simulations of the model under different parameters. Different combinations of learning rates of the adaptive elements of the model (control policy matrix ***Z*** and muscle synergies matrix ***W***), which may have (or not) a nonzero value, are shown in different *columns*. Different performance metrics *(see text*) are shown in different *rows*. Colored *lines* correspond to the three different types of perturbation simulated. Lines and shaded regions indicate average and standard error of each metric across sixteen different model initializations. Shaded rectangles indicate cycles during when each perturbation was applied.

In these simulations, we also examined the ability of the baseline muscle synergies to reconstruct the muscle activity generated by the model over time. Regarding the reconstruction quality (R^2^) of the muscle activity patterns (Fig 3, *third row*), there is a similar decrease for both the rotation and the compatible surgery perturbations, under all three combinations of learning rates. The reconstruction quality R^2^ during the incompatible surgery, however, does not decrease when only the control policy is updated, but it does decrease when the muscle synergies are updated. Statistical analyses (S3 Table) confirm that there is no significant difference in the reconstruction quality R^2^, at the final cycle of perturbation, between the rotation and the compatible surgery perturbations. When only the control policy is updated, there is a statistically significant difference in the R^2^ between the incompatible and the compatible surgeries (*p* = 2.9 ×10^-3^), but not between the incompatible surgery and the rotation (*p* = 0.35); when only the synergies are updated, there are no statistically significant differences between the incompatible surgery and the two other perturbations; and when both control policy and synergies are updated, there is a statistically significant difference between the incompatible surgery and the two other perturbations, suggesting a larger decrease in the R^2^ for the incompatible surgery under that combination of model parameters. A larger decrease in muscle activity reconstruction quality during incompatible surgeries than during compatible surgeries has been observed experimentally [8], although seemingly to a larger degree as what we observe in our simulations.

Regarding the washout period (after the perturbations were removed), we can see a large aftereffect in the force direction error for both the rotation and the compatible surgery, and a comparatively small aftereffect for the incompatible surgery. Aftereffects are considered a hallmark of implicit adaptation and have been reported in a variety of motor tasks [15], and a larger aftereffect for compatible surgeries in comparison with incompatible surgeries has been reported in experiments [8]. Although we can see a large aftereffect in the force magnitude error for the incompatible surgery, there is no published experimental data that we can directly compare this with, since our model concerns the feedforward components of movement while in experiments with virtual surgeries participants were allowed to make online corrections [8,32], making it difficult to discern the magnitude of the feedforward component of the movement. The R^2^ after incompatible surgeries remained at a lower level during the washout than after compatible surgeries. While this has not been observed in [8], in an experiment where participants first performed a task under an incompatible surgery and later under a compatible surgery [10], a persistent decrease in the R^2^ during the second perturbation was observed, which is in line with our simulation results.

We also examined the norm of the muscle activity generated by the model while training with the perturbations (Fig 3, *fourth row*). When both control policy and muscle synergies are updated, we observe a transient increase in the norm of the muscle activity during both the visuomotor rotation and the compatible surgery, mostly at the same time as the force magnitude error is decreasing, with the norm of the muscle activity decreasing slowly after. These transient increase and subsequent slow decrease occur again during the washout. In contrast, during the incompatible surgeries, the increase in muscle activity seems to persist during the perturbation, decreasing only during washout. Statistical analyses (S4 Table) with the data at the last cycle of perturbation indicate statistically significant differences in the norm of the muscle activity for the incompatible surgery compared with the other two perturbations, but no significant differences in the norm of the muscle activity between the rotation and the compatible surgery perturbations, with all combinations of model parameters. While the norm of the muscle activity has not been investigated in previously published data of experiments with virtual surgeries [8,10,32], a quick increase followed by a slow decrease of muscle activity has been observed in arm reaching experiments with velocity-dependent and divergent force fields [33], perturbations which we consider similar to our rotation and compatible surgery since they do not require changes in the structure of the synergies. The slow decrease in the muscle activity after the transient increase is mostly due to the regularization that we implement on the control policy and on the muscle synergies, which will become more evident when we analyze the results of simulation 3, where we compare the models trained with and without regularization.

### 3.3 Subspaces of muscle activity during training

In the previous section, when both the control policy and the muscle synergies in the model were updated, we reported that 1) the reconstruction quality of the muscle activity by the original synergies (R^2^) decreased more during the incompatible surgery and 2) the norm of the muscle activity increased more during the incompatible surgery compared to the two other perturbations. These results suggest differences in the usage of the subspaces of the muscle activity space elicited by the training under the different perturbations. To elucidate how the usage of the muscle activity space changes during the perturbations, we evaluated the norm of the muscle activity projected into different subspaces of the muscle activity space (Fig 4), considering the combination of model parameters when both the control policy and the muscle synergies are updated.

**Fig 4:**
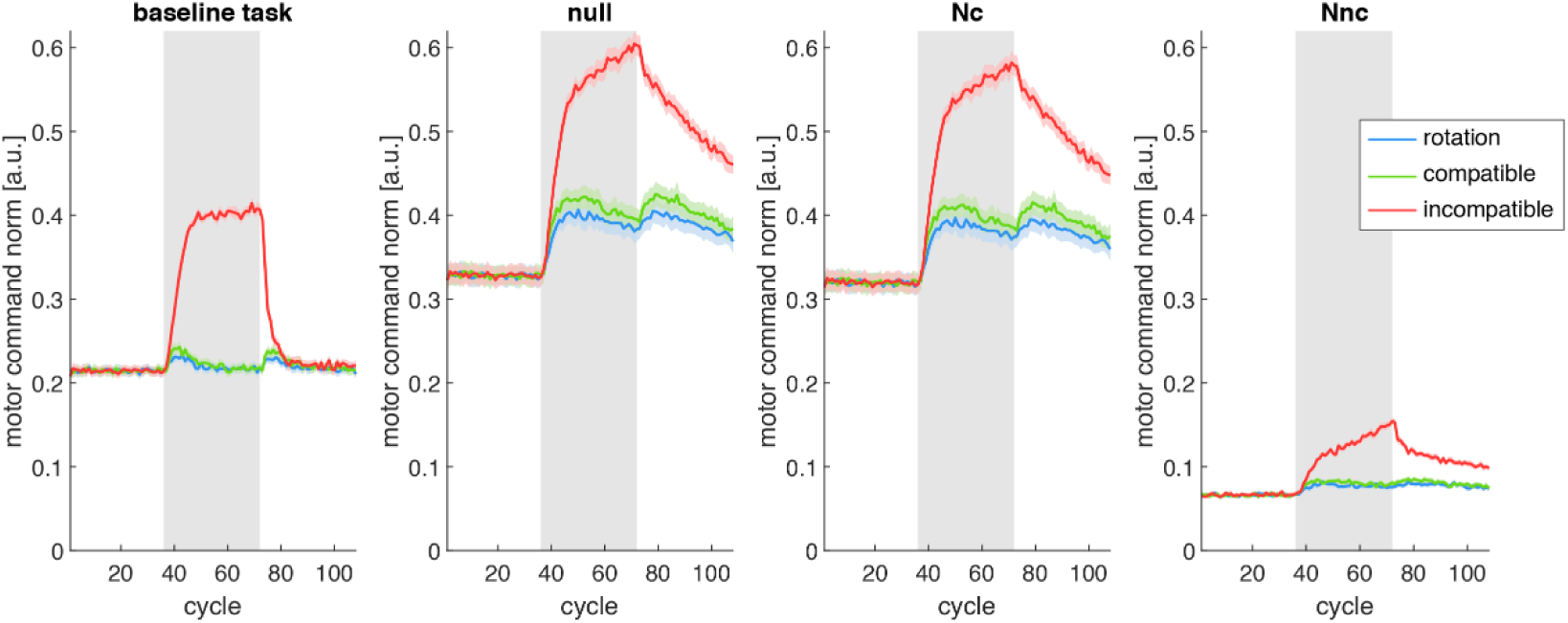
Norm of muscle activity in different subspaces of the muscle activity space. Data from simulation 1, under the model parameter combination with an update of both control policy and muscle synergies. The lines (colored according to the type of perturbation) indicate the norm of the muscle activity projected into different subspaces of the muscle activity space (*columns*), averaged across all targets in a cycle (*gray rectangles* indicate cycles during which each perturbation was applied). Lines and shaded regions are average and standard error across sixteen different model initializations. Null: null space of the baseline task space. Nc: common subspace between null space and baseline synergies. Nnc: subspace of null space not spanned by baseline synergies.

When analyzing the norm of the muscle activity projected in the baseline task space, we can see that it initially increases for all perturbations, however it quickly decreases to near baseline levels for the rotation and the compatible surgery, while remaining large for the incompatible surgery. Statistical analyses at the last perturbation cycle (S5 Table) show a statistically significant difference between the norm of the muscle activity projected in the baseline task space between the incompatible surgery and the two other perturbations, and no significant difference between the rotation and the compatible surgery. This large increase in the norm of the muscle activity projected in the baseline task space for the incompatible surgery is due to an inappropriate forward model at the beginning of the training, trying to correct for movement errors caused by the perturbation (which reduces the span of the baseline synergies to a single dimension) by increasing the patterns of muscle activity in the baseline task space [20], to which it was set during the initialization of the simulation. The increase in the muscle activity decelerates because the forward model appropriate for the incompatible surgery is learned and muscle activity patterns outside the baseline task space are increasingly used. The large norm of the muscle activity in the baseline task space immediately before the removal of the incompatible surgery is what caused the large after-effect in the force magnitude error described in the previous section (Fig 3). After the removal of the perturbation, the norm of the projection of the muscle activity in the baseline task space transiently increases again for both rotation and compatible surgery, while for the incompatible surgery it decreases back to baseline levels (since the environment is the same as the baseline task environment and since the reaching required a specific amount of force, it is expected that the baseline task component of the muscle activity decreases back to baseline levels for all perturbations).

Noticeably, when analyzing the norm of the muscle activity in the null space and in its ***N***_*c*_ (intersection between null space and baseline synergies) and ***N***_*nc*_ (subspace of null space not spanned by baseline synergies) subspaces, we see that most of the muscle activity is in the ***N***_*c*_ space, while the component of the muscle activity in the ***N***_*nc*_ space is much smaller. As in our modular architecture the muscle activity is generated by a combination of synergies, it is expected that, at baseline, most of the muscle activity in the null space can be explained by a combination of muscle synergies (i.e., the ***N***_*c*_ space), while the muscle activity in the null space not spanned by the baseline muscle synergies (i.e., the ***N***_*nc*_ space) is mostly due to motor noise. The increase in the muscle activity in the ***N***_*nc*_ space during the rotation and the compatible surgery is negligible, however during the incompatible surgery it appears comparably larger, which is confirmed by a statistically significant difference in the norm of the muscle activity in the ***N***_*nc*_ space between the incompatible surgery and the two other perturbations (S5 Table). After the removal of the perturbation, the norm of the muscle activity in the ***N***_*nc*_ space slowly decreases for the incompatible surgery, however not fully back to baseline levels until the end of the washout phase.

In summary, the results of the analysis of the muscle activity norm in different subspaces of the muscle activity space show that, during the perturbation, the usage of the subspace of the null space not spanned by the baseline synergies (the ***N***_*nc*_ space) increases more for the incompatible surgery compared with the two other perturbations. These results are in line with the previous results of the reconstruction quality of the muscle activity decreasing more for the incompatible surgery than for the compatible surgery, and show changes in the patterns of muscle activity generated by the model due to the learning process, changes which partially persist for some time even after the removal of the perturbation.

### 3.4 Examining different learning rates of the forward model

In the previous simulation, the forward model of the musculoskeletal system was updated as the models encountered the perturbations. Previous computational models of the trial-by-trial update of motor activity which included a modular architecture have relied on the assumption that the forward model, responsible for the error correction, was known *a priori*, or instantly fully learned as soon as the perturbation is introduced [21,22], which has been criticized as unplausible [27]. In simulation 2, we investigated three possibilities for the learning of the forward model: either no update of the forward model (learning rate of zero), or a slowly updated forward model (non-zero learning rate, the same one used in simulation 1), or the ideal forward model for each perturbation. This allows us to compare the predictions of our model with other models which assumed an ideal forward model and with the available experimental data.

The results of simulation 2 (Fig 5), show that there is no apparent change in any of the metrics during the rotation and the compatible surgery across all three learning rates of the forward model. Statistical analyses on the force direction error (S6 Table), force magnitude error (S7 Table), reconstruction quality of the muscle activity using the original muscle synergies (R^2^) (S8 Table), and norm of the muscle activity (S9 Table) at the final cycle of perturbation confirm that there is no significant difference across any of the parameters for the forward model for both perturbations, in all metrics. For the incompatible surgery, in contrast, there are clearly visible differences in the four metrics across the three learning rates. In comparison to when the forward model is updated slowly, both force direction error and force magnitude error are much higher when the forward model is not updated (even higher than at the onset of the perturbation), and much lower when using the ideal forward model, where the decrease in the error becomes indistinguishable from the two other perturbations (with the exception of the washout, when there is a clear transient increase in the error for the rotation and the compatible surgery which is absent for the incompatible surgery). In terms of the R^2^, there is no significant difference in the decrease between when the forward model is slowly updated and when it is not updated (S8 Table), but there is a significantly larger decrease when the ideal forward model is used, compared to the two other conditions. Compared to when the forward model is updated, the motor command norm when the forward model is not updated reaches a significantly higher level, and a significantly lower level when the ideal forward model is used (S9 Table).

**Fig 5:**
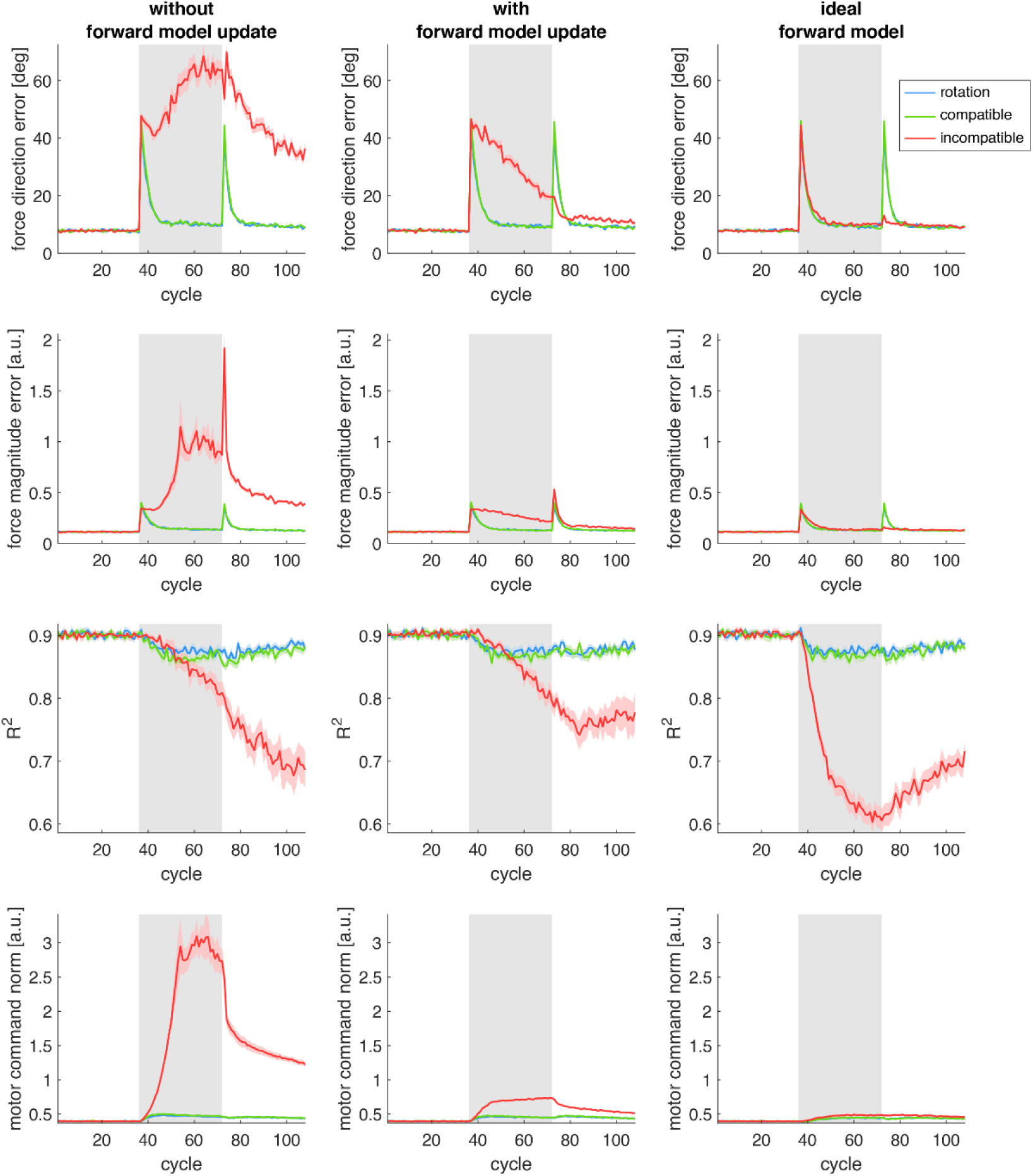
Simulations of the model under different learning rates of the forward model. Three different learning rates of the forward model are presented in the *columns*. Colored lines correspond to the three different perturbations simulated. Lines and shaded regions are average and standard error of each metric (*rows*) across sixteen different model initializations. Shaded rectangles indicate cycles during when each perturbation was applied.

When the forward model is not updated, during the incompatible surgery, there is a large increase in the norm of the muscle activity generated by the model, while the error in the force generated by this muscle activity does not decrease, rather it increases compared to the onset of the perturbation. Since the update of the muscle activity is dictated by the forward model, and since the incompatible surgery requires the usage of novel patterns of muscle activity, the change in the muscle activity generated by the baseline forward model is not effective in the perturbed environment, and the increase in its norm does not reduce the force error.

In contrast, with an ideal forward model, the decrease of the force error during the incompatible surgery is as fast as during the rotation and the compatible virtual surgery, with a large change in the structure of the muscle activity as evidenced by the large decrease in R^2^. Analyzing the norm of the projection of the muscle activity in different subspaces when using the ideal forward model (Fig S1), we see a large increase in the usage of the ***N***_*nc*_ space during the incompatible surgery, and no increase in the norm on the baseline task space. This is in contrast with the condition when the forward model is slowly updated, where the norm of the muscle activity in the baseline task space shows a large increase (Fig 4), and R^2^ does not decrease as much (Fig 5). These results further indicate that, during an incompatible surgery, changes in the baseline task component of the muscle activity can be attributed to an incorrect forward model, and a correct forward model leads to larger changes in the muscle activity, specifically in the ***N***_*nc*_ space, and to a faster decrease of the force error.

### 3.5 Effect of regularization

In the preceding simulations of our model, whenever an element of our model (either the control policy matrix ***Z*** or the muscle synergies matrix ***W***) was updated, we also applied a nonzero regularization to the element, which pulled each element of the matrices towards zero. In simulation 3, we compare two combinations of parameters of the model, with and without regularization, in order to characterize the effect of the regularization. We found that, in general, the regularization has a small but significant effect in both force direction error, force magnitude error, and reconstruction R^2^ (Fig 6), for all perturbations. Statistical analyses on the data at the last cycle of perturbation (S10 Table) suggest a significant decrease of, on average, 1.67° in the force direction error and of 0.024 (arbitrary units) in the force magnitude error, and a significant increase of 0.019 in R^2^ caused by the regularization. A significant and negative effect of the regularization in the motor command norm was also observed (S11 Table), this time with a significant interaction with the perturbation type, with the effect being seemingly stronger for the incompatible surgery (*F(1,90) =* 410.00) than for the compatible surgery (*F(1,90) =* 112.00) and for the rotation (*F(1,90) =* 103.54), possibly because the increase in the norm of the motor commands during the latter two perturbations is not very large.

**Fig 6:**
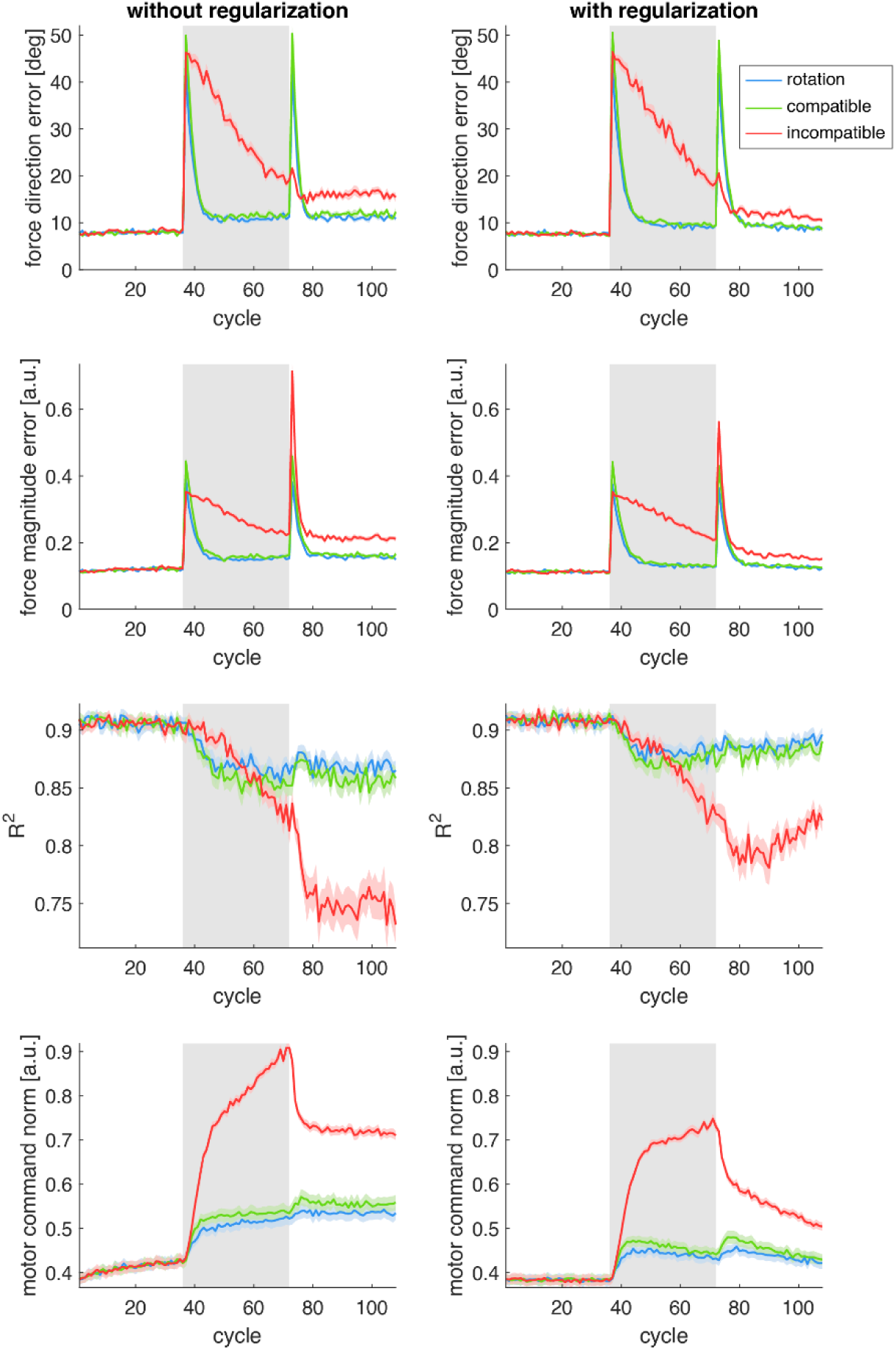
Effect of regularization. Simulations performed with or without regularization are shown in different *columns*. Colored lines correspond to the three different types of perturbation simulated. Lines and shaded regions are average and standard error of each metric (on each *row*) across sixteen different model initializations. Shaded rectangles indicate cycles during when each perturbation was applied.

These results suggest that, while the regularization has a significant effect on the force magnitude error of the model, its largest effect is in decreasing the norm of the muscle activity generated by the model. As there is evidence of the minimization of effort at the muscular [33,34] and at the synergy recruitment level [35], it is then reasonable to include a regularization term in our computational model.

## 4 Discussion

Although the redundancy of the human body affords the central nervous system infinite possibilities in the coordination of multiple joints during movement execution [1], there are strong regularities in the recruitment of muscles [3]. These regularities in the muscle activity patterns can be explained by the existence of a small number of control modules which can be flexibly combined during force generation, simplifying the problem of coordination of the multiple degrees-of-freedom of the musculoskeletal system [2]. A modular architecture for the generation of muscle activity patterns also explains the speed of learning different perturbations of a motor task in terms of whether the perturbation can or cannot be overcome by the original control modules [8]. Computational models that investigated the error-based learning of novel muscle activity patterns did not account for this possible modularity [19,20], and the models that did account for it assumed that the error correction during learning and during perturbations was always ideal, or that the modules were immutable [21,22].

We proposed a modular architecture for the trial-by-trial generation and learning of redundant muscle activity patterns during isometric force-reaching tasks. In our computational model, explicitly defined modules, representing spatial muscle synergies, are recruited by a control policy and are combined for the generation of the muscle activity, which is input into the musculoskeletal system resulting in the generation of a force. The muscle synergies and the control policy can be updated, at different learning rates, with the backpropagation of the force error through a forward model of the musculoskeletal system, which is not assumed to be known *a priori* and must be updated using force prediction error. Our model qualitatively reproduces the main findings of experiments with virtual surgeries that alter the forces generated by the contraction of muscles [8,10], supporting the hypothesis that the generation and learning of muscle activity patterns in humans is done through a modular architecture.

In our computational model, a forward model of the musculoskeletal system is responsible for the correction of movement errors, and this model is updated through force prediction error. During goal-directed movement, the “goal” of the movement is usually defined in terms of a specific posture of the body in space, or the position and orientation of a tool that is being manipulated, however the CNS cannot control these “distal” variables directly, and instead relies on the control of “proximal” variables, such as the muscle activity, in order to realize movement goals. Given an error in the movement (a difference between the executed and the desired movement), a “distal teacher” that transforms a distal error into an appropriate proximal error (a change in the muscle activity) is necessary for the CNS to know how to change its control policy in order to minimize/correct the movement error [16]. It has been argued that the knowledge of this “teacher” responsible for error correction cannot be innate [27], and in novel sensorimotor tasks in which new patterns of muscle activity are required, such as during incompatible virtual surgeries, this internal model for error correction must be learned by the control system in order for the controller to minimize its errors [20]. Indeed, incompatible virtual surgeries could not be learned when the forward model was not updated. Thus, it is reasonable to include in our model a gradual update of the forward model of the musculoskeletal system.

Computational models of trial-by-trial learning that included modularity in the generation of muscle activity have been investigated before. Hagio & Kouzaki (2018) found that modularity may accelerate learning by reducing the bias of the distribution of the mechanical contributions of neurons in the controller neural network. A similar argument has been made by Barradas et al. (2023), where the speed of learning under a perturbed environment was related to the uniformity of the shape of the cost function in that environment, expressed as the ratio between the smallest and the largest eigenvalue of the Hessian matrix of the cost function around its minimum. They showed that, in a non-modular architecture using the distal learning framework, learning after an incompatible surgery would in fact be slightly faster than learning after a compatible surgery, which is explained by their analysis of different uniformities of the cost functions across the two virtual surgeries. Using a modular architecture, they observed that learning after incompatible surgeries was slower than after compatible surgeries. As mentioned previously, one of the key differences between these two models and our model is that we do not assume that the forward model responsible for the error correction is known at the beginning of the perturbation, and must be learned by minimizing the force prediction error.

Another key difference between our model and the model presented in Barradas et al. (2023) is that we assume that the structure of the spatial muscle synergies can be updated by the learning algorithm, while they assumed that the synergies are fixed. It has been hypothesized that changes to the synergy recruitment may be faster than changes to the synergies’ structure [8], which may support their assumption that, for the duration of an experimental session, the synergies are effectively fixed. However, in the simulations without update of the synergies, the reconstruction quality of the muscle activity using the original muscle synergies (R^2^) during the incompatible surgery did not decrease, in contrast with a large decrease in the experimental results with human participants. This indicates that allowing muscle synergies to change over time provides a better fit to the observed experimental data, at the level of changes to the muscle activity.

In our model, when the synergies are not updated, we observed no reduction in the force magnitude error during incompatible surgeries, while in the model of Barradas et al. (2023) there was a slow reduction in the error. This is because, as they explicitly mentioned, they did not utilize the “ground truth” synergies in the calculation of the virtual surgeries, but rather they used an estimate of the synergies from the muscle activity generated by the model during baseline. This may cause the incompatible surgeries computed in this way not to be truly “incompatible” (in the sense that they may still allow the existing synergies to span the entire force space), although this approach has the merit of following more closely the experimental approach of estimating the synergies from the data (because the experimenters do not have access to the ground truth synergies).

The regularization that we implemented in our model, to both control policy and synergy structure matrices, is comparable to the algorithm of weight decay widely used in artificial neural networks to improve their generalization [36]. Weight decay has also been investigated in computational models of sensorimotor transformations [37], where it has been shown that, over tens of thousands of trials of training (and with a regularization weight in the same order of magnitude as we used in our simulations), effort decreases to optimal levels. Our simulations, in contrast, were designed to have a similar number of trials as a typical single-session experiment in humans, which is in the order of hundreds of trials, and therefore were most likely too short for us to observe a complete minimization of effort, although we did observe a significantly lower norm of the muscle activity during the three perturbations when using regularization. In addition, experiments in humans performing three-dimensional isometric forces have shown that, rather the individual muscle activity, the recruitment of muscle synergies seems to be minimized, although sub-optimally [35]. Further experiments are necessary to better elucidate the minimization of effort at the muscular and at the synergistic levels during force generation and during learning.

In summary, our computational model explains the differences in adaptation to different types of virtual surgeries, both at the task and the muscle activity levels, through the update of three elements of the proposed modular control architecture: the recruitment of muscle synergies; the structure of the synergies; and an acquired forward model of the task and motor system. Viewing virtual surgeries as a model for neurorehabilitation, where effective muscle activity patterns must be learned, patient-specific impairments may be related to the update/learning of the different elements in our model and to specific neural substrates. Our model may also be used to explore tailored interventions to enhance recovery from specific impairment.

### 4.1 Limitations

One of the limitations of our model is using a fixed number of muscle synergies. Changes in the number of synergies used for locomotion, due to fractionation and merging, have been reported during development and training [38]. Fractionation and merging of synergies in the affected arm after stroke, compared to the synergies extracted from the unaffected arm, have also been shown [39], indicating that cortical damage may affect the number and the structure of muscle synergies used for arm reaching. Although using a fixed number of synergies might be appropriate to study motor learning processes over a short time interval, such as single-session experiments, a model which includes a variable number of muscle synergies might enable a better understanding of changes in motor coordination that occur throughout development, during long-term skill training, and possibly during motor rehabilitation.

We did not include in our model any cognitive processes, which have been shown to influence adaptation [40]. In a recent experiment where participants had more time to reach the target during the virtual surgeries [32], a larger aftereffect in the direction error of the initial movement was observed for the compatible surgery in comparison with the incompatible surgery, immediately after the removal of the perturbations, which is in line with the predictions of our model regarding the force direction error. These results were accompanied by a larger reaction time during the training under incompatible surgeries, which suggested that explicit strategies were used during the incompatible surgery and that implicit adaptation, resulting in aftereffects [41], was used during the compatible surgery. Although our computational model predicts a decrease in the reconstruction quality of the muscle activity using the original synergies (R^2^), this decrease is smaller compared to what has been observed experimentally [8,32], and its increase during the washout period also appears to be slower than in the experimental data. These results, in addition to the findings of possible cognitive strategies during learning under incompatible surgeries, may suggest that our model lacks a cognitive component which could be responsible for the regulation of motor exploration [42–44].

Finally, our model can only predict the generation of feedforward motor commands and the trial-by-trial learning processes of the muscle activity, while the movements executed by the motor system are generally continuous and learning may occur throughout the movement duration [32,45–48]. We plan to extend our model to also predict the entire time-course of the muscle activity and of the generated force, as well as online feedback error correction mechanisms. A feedback-driven modular architecture in a novel redundant sensorimotor task, including the assumption of no *a priori* knowledge of the sensorimotor task, has been investigated recently [49], however such model did not account for possible changes in the synergies’ structure, only in their recruitment, and did not include a feedforward component. In a recent virtual surgeries experiment, it has been shown that increasing the trial duration leads to a faster improvement in participants’ online feedback error correction compared to their initial movement error (related to trial-by-trial changes in the feedforward generation of the movement) during incompatible surgeries [32]. This faster improvement in the online feedback error correction, attributed to the longer movement durations leading to more exploration of the task, suggests a possible dissociation between feedback and feedforward error correction mechanisms, which has also been observed in mirror-reversal experiments [50]. In terms of modeling, this might suggest different gains for the error correction and the feedforward and the feedback components of the model, or possibly that the trial-by-trial feedforward error correction and the online feedback error correction are made (and learned) through separate mechanisms, each with its own learning rate. Expanding our model to generate the entire movement trajectory could allow us to better understand these observed phenomena.

## Acknowledgments

AD gratefully acknowledges the funding provided by the project “HARIA - Human-Robot Sensorimotor Augmentation - Wearable Sensorimotor Interfaces and Supernumerary Robotic Limbs for Humans with Upper-limb Disabilities” (EU Horizon Europe, GA No. 101070292, https://clem.diism.unisi.it/~haria/). DB gratefully acknowledges the funding provided by the Italian Ministry of Health (IRCCS Fondazione Santa Lucia, Ricerca corrente; GR-2019-12370271,

https://www.salute.gov.it/portale/bandiGara/dettaglioArchivioBandiGara.jsp?lingua=italiano&id=208). The funders had no role in study design, data collection and analysis, decision to publish, or preparation of the manuscript.

## Supporting Information

**S1 Text: Derivations of update equations of the computational model**

**S1 Fig: Norm of muscle activity in different subspaces of the muscle activity space.** Data from simulation 2, with the models using an ideal forward model during each perturbation. The lines (colored according to the type of perturbation) indicate the norm of the muscle activity projected into different subspaces of the muscle activity space (*column*s), averaged across all targets in a cycle (gray rectangles indicate cycles during which each perturbation was applied). Lines and shaded regions are average and standard error across sixteen different model initializations. Null: null space of the baseline task space. Nc: common subspace between null space and baseline synergies. Nnc: subspace of null space not spanned by baseline synergies.

**S1 Table: results of statistical tests on the model with force direction error as the dependent variable, on data from simulation 1.** For the post-hoc comparisons, p-values were Bonferroni-corrected. Comp: compatible (surgery); incomp: incompatible (surgery); rot: rotation.

**S2 Table: results of statistical tests on the model with force magnitude error as the dependent variable, on data from simulation 1.** For the post-hoc comparisons, p-values were Bonferroni-corrected. Comp: compatible (surgery); incomp: incompatible (surgery); rot: rotation.

**S3 Table: results of statistical tests on the model with the reconstruction quality (*R***^**2**^**) of the muscle activity using the original muscle synergies as the dependent variable, on data from simulation 1.** For the post-hoc comparisons, p-values were Bonferroni-corrected. Comp: compatible (surgery); incomp: incompatible (surgery); rot: rotation.

**S4 Table: results of statistical tests on the model with the motor command norm as the dependent variable, on data from simulation 1.** For the post-hoc comparisons, p-values were Bonferroni-corrected. Comp: compatible (surgery); incomp: incompatible (surgery); rot: rotation.

**S5 Table: results of statistical tests on the models with the motor command norm in different subspaces as the dependent variable, on data from simulation 1.** For the post-hoc comparisons, p-values were Bonferroni-corrected. Comp: compatible (surgery); incomp: incompatible (surgery); rot: rotation.

**S6 Table: results of statistical tests on the model with force direction error as the dependent variable, on data from simulation 2.** For the post-hoc comparisons, p-values were Bonferroni-corrected. Update: update of forward model; no update: without update of forward model; ideal: ideal forward model.

**S7 Table: results of statistical tests on the model with force magnitude error as the dependent variable, on data from simulation 2.** For the post-hoc comparisons, p-values were Bonferroni-corrected. Update: update of forward model; no update: without update of forward model; ideal: ideal forward model.

**S8 Table: results of statistical tests on the model with the reconstruction quality (*R***^**2**^**) of the muscle activity using the original muscle synergies as the dependent variable, on data from simulation 2.** For the post-hoc comparisons, p-values were Bonferroni-corrected. Update: update of forward model; no update: without update of forward model; ideal: ideal forward model.

**S9 Table: results of statistical tests on the model with the motor command norm as the dependent variable, on data from simulation 2.** For the post-hoc comparisons, p-values were Bonferroni-corrected. Update: update of forward model; no update: without update of forward model; ideal: ideal forward model.

**S10 Table: results of statistical tests on the models with the force direction error, the force magnitude error, or the muscle activity reconstruction quality (*R*^2^) using the original synergies as dependent variable, on data from simulation 3.**

**S11 Table: results of statistical tests on the model with the motor command norm as the dependent variable, on data from simulation 3.** For the post-hoc comparisons, p-values were Bonferroni-corrected. Reg: regularization; no reg: without regularization.

